# The *Medicago truncatula* LYR4 intracellular domain serves as a scaffold in immunity signaling independent of its phosphorylation activity

**DOI:** 10.1101/2024.10.23.619817

**Authors:** Bine Simonsen, Henriette Rübsam, Marie Vogel Kolte, Maria Meisner Larsen, Christina Krönauer, Kira Gysel, Mette Laursen, Feng Feng, Gülendam Kaya, Giles E. D. Oldroyd, Jens Stougaard, Sébastien Fort, Simona Radutoiu, Kasper Røjkjær Andersen

## Abstract

Plants perceive and respond to chitin derived from fungal cell walls through lysine motif (LysM) receptor kinases. In the model legume *Medicago truncatula*, CERK1 and LYR4 represent the LysM receptor pair important for chitin-triggered immunity signaling. Here, we show that both the active kinase receptor CERK1 and the pseudokinase receptor LYR4 contribute to immunity signaling, leading to the production of reactive oxygen species (ROS). We determine the crystal structure of the LYR4 core intracellular domain with a bound nucleotide analog in the active site. Biochemical characterization shows that LYR4 binds ATP and has both autophosphorylation as well as transphosphorylation activity towards CERK1. However, *in planta* experiments demonstrate that the phosphorylation ability is not necessary for the function of LYR4 in chitin-triggered ROS production, but that the presence of its intracellular domain is indispensable. Together, we show that in chitin-triggered immunity the intracellular domain of LYR4 serves as a signaling scaffold independent of its catalytic activity.

## Introduction

Plants engage in a wealth of interactions with beneficial and pathogenic bacteria and fungi, and therefore need to monitor their surroundings. To this end, plant cell-surface receptors, such as lysine motif (LysM) receptors, perceive microbe-associated molecular patterns (MAMPs), which can both elicit immunity responses and initiate the establishment of symbiotic associations^1^. LysM receptors have an ectodomain built of three interconnected LysM domains and most feature a single transmembrane helix and either an active kinase or a pseudokinase intracellular domain^2^. The main ligands bound by LysM receptors are carbohydrates containing *N*-acetylglucosamine moieties, namely chitooligosaccharides (CO; chitin), lipochitooligosaccharides (LCO) and peptidoglycans^1–6^.

In *Arabidopsis thaliana* (*Arabidopsis*), the LysM receptor-like kinase (RLK) CERK1 is indispensable for chitin-triggered immunity, while the pseudokinase receptors LYK4 and LYK5 differentially contribute to defense signaling^7–11^. The *lyk4 lyk5* double mutant shows a complete loss of reactive oxygen species (ROS) production, whereas inconsistant results have been reported on the phenotypes of the single mutants and on the interaction of CERK1 with LYK4 and LYK5^9–11^. A recent study showed that both LYK4 and LYK5 form a complex with CERK1 upon chitin binding and contribute to MAPK activation, yet that LYK5 primarily mediates phosphorylation of CERK1, whereas LYK4 is crucial for ROS production^12^.

In the model legume *Lotus japonicus* (*Lotus*), the LysM RLKs CERK6 (previously called LYS6) and the tandem duplicated receptors LYS13 and LYS14 are involved in chitin-triggered immunity^4^. LYS13 and LYS14 are classified pseudokinases and their expression has been shown to increase in root and shoot upon chitin treatment of the plant^13,14^. Single mutants of *lys13* and *lys14* respond to chitin similarly to wild type, suggesting functional redundancy between LYS13 and LYS14, whereas *cerk6* is unable to produce ROS upon chitin treatment and upregulation of defense-response genes and phosphorylation of MAPK3/6 is impaired^4^. Further investigation of chitin triggered immunity signaling in *Lotus* is difficult, as a *lys13 lys14* double mutant is not available.

In the model legume *Medicago truncatula* (*Medicago*), CERK1 (previously called LYK9) is a homologue of *Lotus* CERK6 and LYR4 is a homologue of *Lotus* LYS13 and *Lotus* LYS14^1,4,15^. *Medicago* CERK1 (CERK1 hereafter) and *Medicago* LYR4 (LYR4 hereafter) have been reported to be indispensable for chitin-triggered immunity with both *cerk1* and *lyr4* mutants having an increased lesion size upon infection of leaves with *Botrytis cinerea*^4^. In addition, ROS responses were absent and phosphorylation of MAPK3 and MAPK6 decreased upon chitin octamer (CO8) treatment in both mutants compared to wild type^4,16^.

CERK1 has an active kinase intracellular domain containing all canonical motifs of a eukaryotic protein kinase, while LYR4 is a classified pseudokinase due to the degeneration of several key kinase motifs: a truncated glycine-rich loop, the DFG-motif is NFG, and the HRD-motif is HKN^1,14^ (Figure 1A). Canonically, the glycine-rich loop and the regulatory lysine on the β3-strand stabilize phosphates of bound ATP. Aspartate from the DFG-motif is important for positioning of phosphates for phosphate transfer and the arginine and the aspartate from the HRD-motif contribute to ordering of the activation loop and activation of the incoming substrate, respectively^17,18^. In this study, we investigate how the pseudokinase domain of the *Medicago* LYR4 LysM receptor kinase mediates downstream signaling in immunity. We determine the crystal structure of the LYR4 intracellular domain with an ATP-analogue bound in a non-canonical manner and show that it is catalytically active despite its lack of canonical kinase features. Biochemical characterization and *in planta* ROS production assays of LYR4 variants show that LYR4 plays a scaffolding role in chitin-triggered signaling independent of ATP binding and kinase activity.

**Figure 1.**
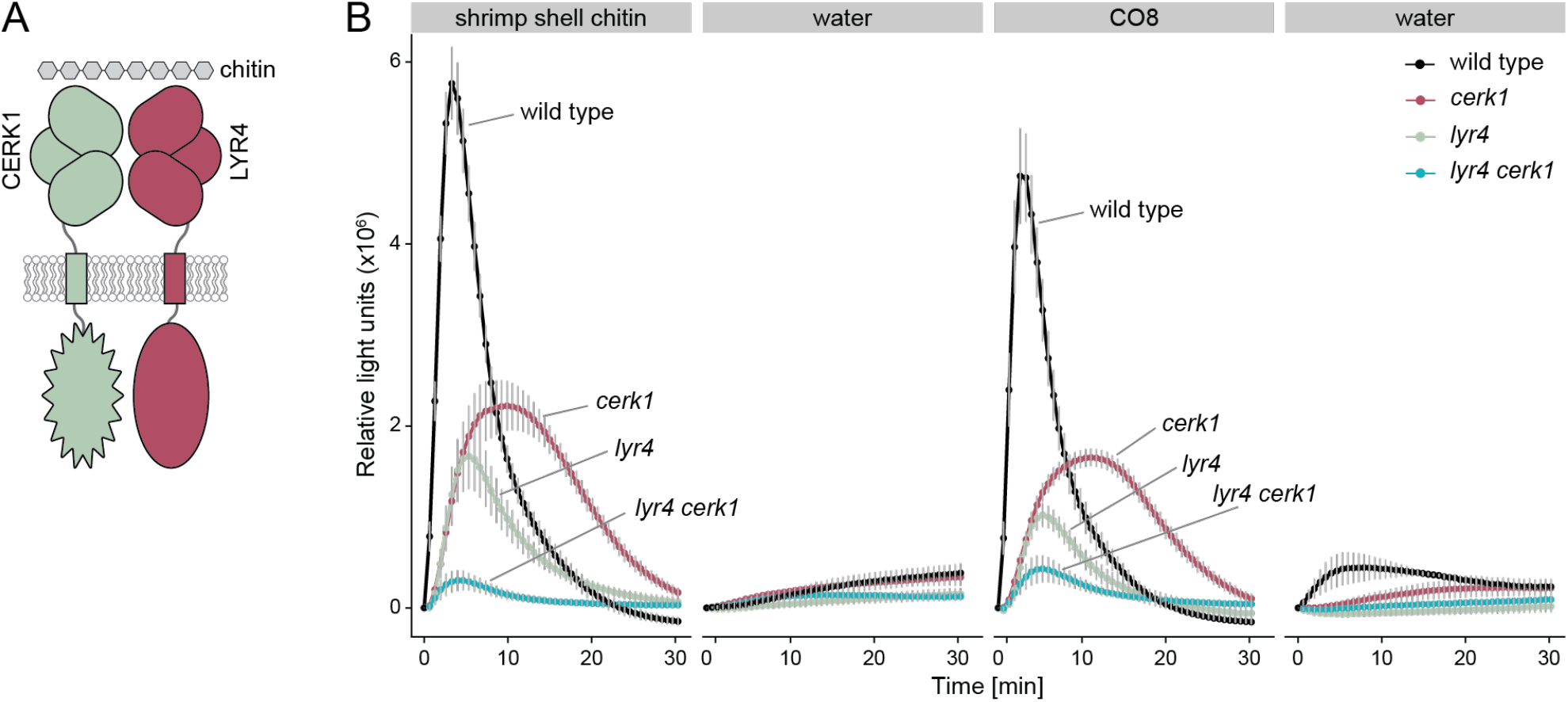
The production of reactive oxygen species in response to chitin is attenuated in *lyr4* and *cerk1*. **A**. Schematic of the *Medicago* LysM-receptors CERK1 and LYR4 in the plasma membrane recognizing chitin. CERK1 is depicted with an active kinase domain and LYR4 with a pseudokinase domain. **B**. Reactive oxygen species produced in wild type, *cerk1, lyr4*, and *cerk1 lyr4 Medicago* roots over a period of 30 minutes after treatment with 0.1 mg/mL shrimp shell chitin, 1 μM chitooctaose (CO8) or water (mean ± standard error of the mean, n = 8 (chitin, CO8) or n = 4 (water)).

## Results

### *Medicago* CERK1 and LYR4 can contribute individually to signaling in immunity

To investigate the individual contributions of CERK1 and LYR4 in immunity signaling, we tested *Medicago cerk1* and *lyr4* single mutants and a *cerk1 lyr4* double mutant for their ability to elicit a ROS response when treated with shrimp-shell chitin or purified CO8 (Figure 1B). While we see almost no ROS response in the *cerk1 lyr4* double mutant, both *cerk1* and *lyr4* single mutants elicit a ROS response when treated with either shrimp-shell chitin or purified CO8. ROS production in both single mutants is, however, attenuated compared to wild type and the ROS curve is broadened in *cerk1* and peaks later than wild type. From this we conclude that ROS production in response to chitin does not fully depend on the presence of both receptors, but that CERK1 and LYR4 can also individually contribute to immunity signaling.

### Crystal structure of LYR4 kinase reveals nucleotide binding ability

To understand how LYR4 mediates signaling, we purified and crystallized its intracellular domain truncated of the juxtamembrane part and the predicted flexible C-terminal tail (Figure S1A). We obtained crystals in a condition supplemented with the ATP analogue adenylyl-imidodiphosphate (AMP-PNP). Diffraction data were collected and resulted in a 3.1 Å resolution structure of the LYR4 kinase processed anisotropically to 2.1 Å (PDB-ID 8PS7) (Figure 2A, Table 1). Based on its amino acid sequence, LYR4 is defined as a pseudokinase due to a truncated glycine-rich loop and degenerated DFG- and HRD-motifs. The electron density revealed a clearly defined AMP-PNP in the nucleotide binding site (Figure 2B), showing that LYR4 kinase has retained nucleotide binding abilities. Closer examination of the binding site revealed that LYR4 binds AMP-PNP in a non-canonical way that compensates for its lack of traditional binding motifs (Figure S1B). Pseudokinases can display compensatory mechanisms to retain nucleotide-binding despite degraded motifs, which has been proposed as an important feature for pseudokinase function^19–21^. Canonically, the ATP phosphates are coordinated by the glycine-rich loop and the regulatory lysine, and while LYR4 does indeed employ the regulatory lysine (K379) to coordinate the α-phosphate, the glycine-rich loop is too short to reach the binding site. Instead, N494 from the modified DFG-motif coordinates both the α- and the β-phosphate while R514 from the activation loop coordinates both the β- and the γ-phosphate.

**Figure 2.**
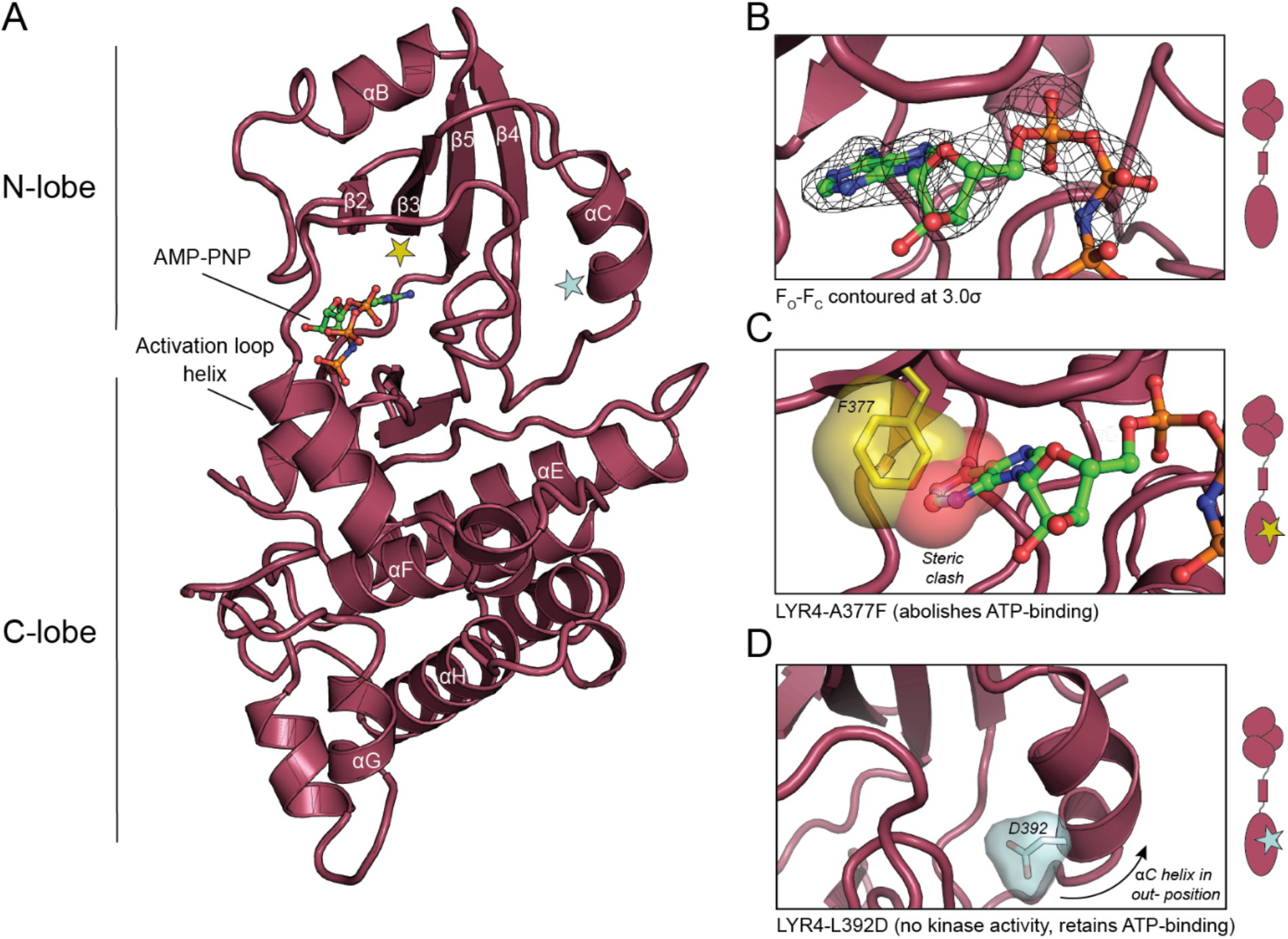
Crystal structure of the LYR4 kinase with a bound ATP analogue. **A**. Crystal structure of the LYR4 kinase bound to the ATP-analogue AMP-PNP annotated with conserved secondary structural elements. Stars in yellow and blue indicate positions of the variants described in C and D. **B**. Omit map of AMP-PNP bound in the nucleotide binding site of LYR4 kinase. The F_O_-F_C_ electron density map is contoured at 3.0σ. **C**. Structural model of LYR4-A377F, where nucleotide binding is obstructed as the phenylalanine (yellow) clashes sterically with the nucleotide (red). **D**. Structural model of LYR4-L392D, which abolishes kinase activity but maintains nucleotide binding as the charged aspartate (blue) pushes the αC in an out-position and breaks the R-spine.

Active kinases have two hydrophobic spines going through the kinase core – the regulatory (R-) spine and the catalytic (C-) spine. In the reaction cycle, kinases assemble and break these spines as they toggle between active and inactive state. The C-spine is assembled when the kinase binds a nucleotide, while the R-spine is assembled when the αC helix moves to an in-position and forms a salt bridge to the regulatory lysine. Both spines need to be assembled for a kinase to be able to phosphorylate its substrate^17,18,22^. LYR4 crystallized in an inactive conformation, as it has an intact C-spine, but a broken R-spine due to the αC helix being in an out-position (Figure S1C, D). Furthermore, the activation loop shows a defined helix in the C-terminus (Figure 2A, labelled activation loop helix). This helix blocks substrate access to the active site while also bringing R514 into position for coordinating the β- and the γ-phosphate of AMP-PNP, representing an yet undescribed inactive kinase conformation. Guided by the structure, we designed two receptor variants to investigate whether nucleotide binding is important for the signaling function of LYR4. LYR4-A377F has a phenylalanine in the nucleotide binding site, which will abolish ATP-binding (Figure 2C). LYR4-L392D introduces a negative charge on the αC helix, making it unable to move to its active in-position and assemble the R-spine, which will keep the kinase inactive while retaining ATP binding (Figure 2D).

### LYR4 serves a scaffold function in immunity independent of its catalytic activity

We expressed and purified the kinase domains of wild type LYR4 and the variants LYR4-A377F and LYR4-L392D (Figure S2) and determined their ability to bind ATP by testing their thermostability using nano differential scanning fluorimetry (nano-DSF) with and without ATP and magnesium (Figure 3A). ATP together with magnesium increased the stability of LYR4 by 4.1°C, indicating that LYR4 binds ATP. While LYR4-L392D was likewise stabilized by 3.7°C, when ATP and magnesium were present, LYR4-A377F was only marginally stabilized by 1.5°C, indicating no, or very weak, ATP-binding. To test LYR4 for potential kinase activity, we performed *in vitro* kinase assays using ^32^P-labeled ATP. We observed that LYR4 can both autophosphorylate and transphosphorylate myelin basic protein (MBP), making it an active pseudokinase (Figure 3B, lanes 3-4). As expected, neither LYR4-L392D nor LYR4-A377F showed kinase activity (Figure S3). CERK1 has autophosphorylation activity, but the *in vitro* activity is low compared to LYR4 (Figure 3B, lanes 3 and 5). Next, we investigated the ability of LYR4 and CERK1 to transphosphorylate each other. We created a version of CERK1 that lacked the catalytic lysine (CERK1-K349A) (Figure S2), making it an inactive kinase (Figure 3B, lanes 9–10), and used it to show that LYR4 can transphosphorylate CERK1 (Figure 3B, lane 11). Conversely, we used the kinase-inactive LYR4-L392D and showed that CERK1 does not transphosphorylate LYR4 *in vitro* (Figure 3B, lane 12).

**Figure 3.**
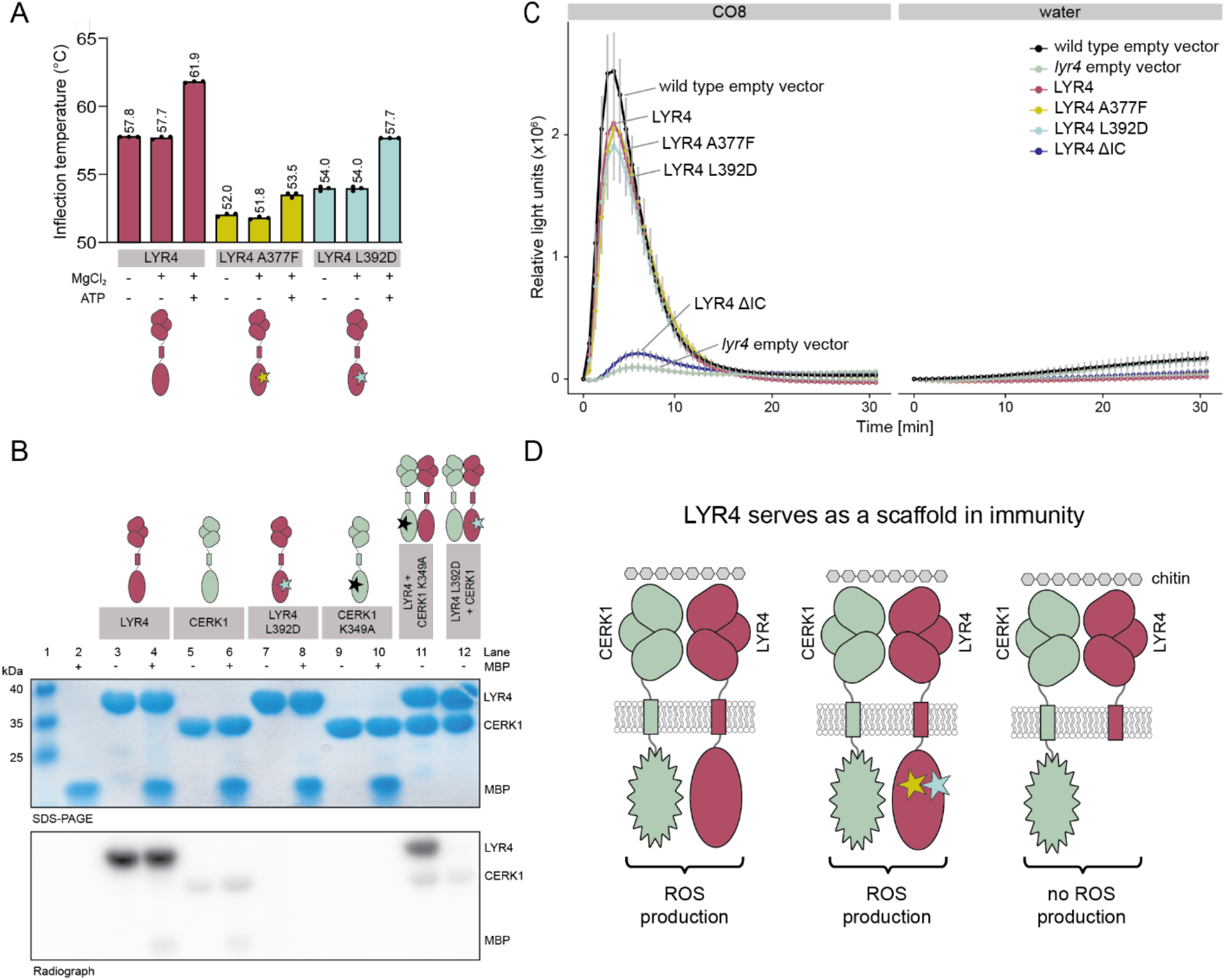
LYR4 functions in immunity signaling independent of ATP-binding and kinase activity. **A**. Nano-DSF thermal stability assay of intracellular domain of LYR4 (bordeaux), LYR4-A377F (yellow) and LYR4-L392D (blue) with and without 1 mM MgCl_2_ and 1 mM ATP. All data points are shown (n = 3). **B**. *In vitro* kinase assay of LYR4, CERK1, LYR4-L392D, CERK1-K349A, and combinations of these incubated with radioactive ATP in the presence and absence of myelin basic protein (MBP) as a transphosphorylation target. Top: Coomassie blue stained SDS-PAGE. Bottom: Corresponding radiograph. **C**. ROS production in *Medicago* wild type and *lyr4* single mutant roots transformed with *Lyr4, Lyr4*-A377F, *Lyr4*-L392D or *Lyr4*-ΔIC over a period of 30 minutes after treatment with 1 μM CO8 or water (mean ± standard error of the mean, n = 15 (CO8) or n = 3 (water)). **D**. Model of chitin-induced ROS production mediated by *Medicago* LYR4. Neither phosphorylation activity nor nucleotide binding by LYR4 are needed to mediate an immunity response to chitin (A377F and L392D mutations represented by yellow and blue stars, respectively), but presence of the LYR4 intracellular domain is crucial, as it serves as a scaffold for complex formation or downstream interacting partners.

Since LYR4 binds ATP and has phosphotransfer activity, we wondered whether these abilities impact LYR4 signaling function in immunity. To test whether LYR4 signaling is dependent on either kinase activity or nucleotide binding and subsequent conformational changes, we transformed *Medicago lyr4* plants with native *Lyr4, Lyr4-A377F* (no nucleotide binding), or *Lyr4-L392D* (kinase inactive, but retained nucleotide binding). We then tested transformed roots for their ability to elicit ROS upon treatment with CO8. Plants expressing *Lyr4, Lyr4-A377F*, or *Lyr4-L392D* all produced ROS equal to the response in wild type *Medicago* transformed with an empty vector control when treated with CO8 (Figure 3C), showing that neither the LYR4 kinase activity nor nucleotide binding is important for its ability to signal in chitin-triggered immunity.

To test whether the kinase domain is required at all, we constructed a LYR4 variant lacking the entire intracellular domain, *Lyr4-ΔIC*, and tested it for its ability to complement LYR4 in *Medicago lyr4* hairy roots. When expressed in *Nicotiana benthamiana*, LYR4-ΔIC can be trafficked to the plasma membrane indicating that the protein is produced and properly folded (Figure S4A). LYR4-ΔIC could, however, not mediate a ROS response in *Medicago lyr4* hairy root experiments (Figure 3C, S4B), showing that the intracellular domain of LYR4 is crucial for immune signaling. Together, our results demonstrate that the intracellular part of LYR4 serves as a scaffold in chitin-mediated immunity signaling independent of its ATP binding and phosphorylation activity.

## Discussion

Pseudokinases comprise up to 17% of the plant kinome and are crucial for signaling in numerous pathways, for example those dependent on LysM receptors^23^. However, the understanding of pseudokinase signaling mechanisms in plants remains limited. In this study, we investigate the function of *Medicago* LYR4 in chitin-triggered immunity.

We show that LYR4 can function in perception of chitin in the absence of CERK1, although ROS levels are attenuated. Previously, both the *cerk1* and the *lyr4* mutants were shown to completely lack a ROS response upon CO8 treatment^4,16^. The lower ROS response in *cerk1* compared to wild type could be due to LYR4 pairing with a less efficient or less abundant signaling partner. CERK1 is part of the LYK subfamily of LysM-RLKs harboring potentially active kinases, which has 11 members in *Medicago*, while LYR4 is part of the LYR subfamily that also has 11 members in *Medicago*, with LYR7 being the phylogenetically closest^1^. Some of these less described LysM receptors might be able to work together with CEKR1 and LYR4 in chitin-triggered immunity.

To better understand signaling mediated by a pseudokinase, we determined the crystal structure of the LYR4 kinase with a bound ATP-analogue. The nucleotide was clearly defined in the electron density, which prompted us to investigate the functional relevance of nucleotide binding and phosphotransfer ability. Thermostability assays additionally confirmed that LYR4 kinase binds ATP and closer inspection of the structure showed that the ATP-analogue is bound in a non-canonical manner. Despite lacking canonical binding motifs, several pseudokinases have demonstrated the ability to bind nucleotides through various mechanisms and the binding event itself has been proposed to lead to a conformational regulation^20,21,24^. *In vitro* kinase activity assays showed that LYR4 has both auto- and transphosphorylation activity making it an active pseudokinase. In general, many classified pseudokinases are active when tested experimentally, as they make up for lacking motifs by employing modified catalytic mechanisms^21,25^.

We investigated whether LYR4 is dependent on conformational changes prompted by nucleotide-binding or on kinase activity for signaling in immunity. Complementation studies in hairy roots demonstrated that neither ATP-binding nor kinase activity are crucial for LYR4 signaling. It is intriguing that LYR4 kinase has retained nucleotide binding ability despite the apparent lack of functional relevance, but we cannot exclude the possibility that nucleotide binding plays a role in other signaling pathways, such as AM symbiosis. Here, we show that the presence of the LYR4 intracellular domain is indispensable for full ROS elicitation upon chitin treatment (Figure 3D). We conclude that LYR4 serves a non-catalytic scaffold function in immunity signaling and that CERK1 kinase activity is responsible for signaling downstream of the putative CERK1-LYR4 receptor complex. Similarly, presence of the kinase domain of LYK5, the closest homologue to LYR4 in *Arabidopsis*, is essential for chitin signaling, even though the kinase has no phosphorylation activity^9^. In a broader view, scaffolding pseudokinase receptors like LYR4 and *Arabidopsis* LYK5 might be necessary to correctly arrange their co-receptors CERK1, as well as mediate interactions with downstream signaling partners. Future studies will hopefully elucidate how kinase and pseudokinase receptor complexes are formed to mediate signaling.

## Acknowledgments

We thank Majken Kiel Sørensen for plant care and seed production at our greenhouse, Sara Bährentz for help with plant work, and Leila Margot Henkes for proofreading the manuscript. We acknowledge the Diamond Light Source for provision of experimental facilities and the staff of beamline I04 for assistance with data collection. This work was funded by the Novo Nordisk Foundation (NNF18OC0052855), the Danish Council for Independent Research (3103-00137B), the Carlsberg Foundation (CF21-0139), the Agence Nationale de la Recherche through Labex ARCANE and CBH-EUR-GS (ANR-17-EURE-0003), Glyco@Alps (ANR-15-IDEX-02), ICMG (UAR 2607) Grenoble Chemistry Nanobio mass spectrometry platform, the USDA-NIFA grant (2022-67014-38607), and the project Enabling Nutrient Symbioses in Agriculture (ENSA), that is funded by Bill & Melinda Gates Agricultural Innovations (INV-57461), the Bill & Melinda Gates Foundation and the Foreign, Commonwealth and Development Office (INV-55767).

## Author contributions

Investigation – BS, HR, MVK, MML, CK, GK. Formal analysis – BS, HR, KG, ML, JS, SR, KRA. Resources – SF, FF. Visualization – BS, HR. Conceptualization – SR, KRA. Supervision – KG, SR, KRA. Project administration – KRA. Funding acquisition – GEDO, SR, KRA. Writing – original draft preparation – BS, HR, KRA. Writing – review & editing – All authors.

## Data availability

The coordinates and structure factors for the crystal structure of the LYR4 kinase domain have been deposited in the Protein Data Bank (PDB-ID 8PS7).

## Materials and Methods

### Plant material

*Medicago truncatula* cv. R108 was used as wild type. The mutants *lyr4-2* (NF15280) and *cerk1* (NF16753) were reported previously^4,26^. The double mutant *cerk1 lyr4-*1 was generated by crossing *cerk1* and *lyr4-1*(NF10265) plants. The *lyr4-1* and *lyr4-2* mutants exhibit similiar responses to CO8 treatment as previously described^4^.

### Bacteria and culture conditions

*Escherichia coli* TOP10 (Thermo Fisher Scientific) were used for molecular cloning and grown in LB medium at 37°C. Chemically competent *E. coli* Rosetta 2 cells (Sigma Aldrich) were used for protein expression and grown as specified below. *Agrobacterium rhizogenes* strain AR1193 was used for hairy root transformations and *Agrobacterium tumefaciens* strain AGL1 was used for transient transformation of *Nicotiana benthamiana. Agrobacterium* strains were grown in LB medium at 28°C.

### Medicago stable mutant line ROS assay

*Medicago* wild type, *cerk1, lyr4* and *cerk1 lyr4* seeds were treated with sulfuric acid for 3 min, washed with ddH_2_O, surface sterilized with 3% sodium hypochlorite, washed again with ddH_2_O and incubated in ddH_2_O for two hours. Swollen seeds were transferred to square plates with wet filter paper and incubated at 21°C under 16/8-h light/dark conditions for 2 days. Seedlings were then transferred to square plates with a slope of ¼ B&D medium^27^ solidified with 1.4% Agar Noble (Difco). The agar was covered with wet filter paper (AGF 651; Frisenette ApS) and a metal bar with ten holes for roots was placed on top of the slope. Plates were kept in boxes excluding light from the roots below the metal bars. After three days on plates, seedlings were cut below the hypocotyl, and one full root was transferred to each well of a white 96-well flat-bottomed polystyrene plate (Greiner Bio-One) and kept overnight in sterile water. The water was then replaced by a reaction mixture consisting of 0.5 mM L-012 (Wako Chemicals), 5 μg/mL horseradish peroxidase (Sigma Aldrich), and 1 μM octa-N-acetyl-chitooctaose (CO8, obtained from Sébastien Fort^28^) or 0.1 mg/mL shrimp-cell chitin (Sigma Aldrich). Luminescence was recorded with a Varioskan LUX multimode microplate reader (Thermo Fisher Scientific) for 30 minutes.

### Generation of plant expression vectors

For generation of *Medicago truncatula* expression vectors, sequences of *LYR4* and *LYR4* variants were assembled with the native *LYR4* promoter and the 35S terminator via Golden Gate cloning^29^. The constructs were thereafter cloned into the pIV10 expression vector alongside a construct containing three YFP fluorophores fused to a nuclear localization signal (tYFP-NLS), which serves as a transformation marker. For a localization experiment in *N. benthamiana*, a sequence was assembled in a pICH vector^29^. An overview of protein sequences used for constructs is provided in Additional Data 1.

### Hairy root transformation

*Medicago* wild type and *lyr4* seeds were treated with sulfuric acid for 3 min, washed with ddH_2_O, surface sterilized with 3% sodium hypochlorite, washed again with ddH_2_O and incubated in ddH_2_O for two hours. Swollen seeds were transferred to square plates with wet filter paper and incubated at 21°C under 16/8-h light/dark conditions for 2 days. Seedlings were then transferred to square plates with a slope of agar solidified with 0.8% Gelrite (Duchefa Biochemie) and supplemented with ½ Gamborg’s B5 nutrient solution (Duchefa Biochemie). Transconjugant *A. rhizogenes* AR1193 strains carrying the construct of interest were grown for three days on LB Agar with Ampicillin, Rifampicin and Spectinomycin, each at a final concentration of 100 μg/mL. Cells were resuspended in 4 mL YMB (5 g/L mannitol, 0.5 g/L yeast extract, 0.5 g/L K_2_HPO_4_, 0.2 g/L MgSO_4_ x 7 H_2_O, 0.1 g NaCl, pH = 6.8). 4-day-old seedlings were then transformed using a 1 mL syringe with a needle (Sterican Ø 0.40×20 mm), wounding the hypocotyl and placing a droplet of the bacterial suspension on the wound. The plates with the transformed seedlings were kept in the dark for one day and then at 21°C under 16/8-h light/dark conditions. After three weeks, primary roots were removed and seedlings with emerging transformed roots were transferred to Magenta boxes (Sigma-Aldrich) filled with clay aggregate (LECA, 2-4 mm; Saint-Gobain Weber A/S) supplemented with 80 mL 1/4x B&D nutrient solution with 3 mM KNO_3_.

### ROS assay in Medicago hairy roots

Variants of LYR4 were tested for their ability to complement the *lyr4* mutant. After three weeks of growth in Magenta boxes (Sigma-Aldrich), hairy roots expressing the tYFP-NLS transformation marker were cut into small pieces of approximately 1 mm in length. Pieces from about 1.5 cm of total root length from one plant were transferred into a single well of a white 96-well flat-bottomed polystyrene plate (Greiner Bio-One) containing sterile water. The root cuttings were left in water at room temperature in the dark overnight. Afterwards, the water was removed and a reaction mixture containing 0.5 mM L-012 (Wako Chemicals), 5 μg/mL horseradish peroxidase (Sigma), and 1 μM octa-N-acetyl-chitooctaose (CO8, obtained from Sébastien Fort^28^) or water was added. Luminescence was recorded with a Varioskan LUX multimode microplate reader (Thermo Fisher Scientific) for 30 minutes. Values derived from curves without a single clear peak within 30 min were excluded.

### Transient transformation of *N*. *benthamiana*

Agrobacterium mediated transformation of *N. benthamiana* leaves was performed as previously described^30^. In short: *A. tumefaciens* AGL1 bacteria carrying the *Lyr4*-ΔIC-*GFP* construct driven by the *LYR4* promoter were resuspended in infiltration solution (10 mM MgCl_2_, 10 mM MES, 150 μM acetosyringone, pH 5.6) at an OD_600_=0.025, incubated for two hours in the dark and infiltrated into leaves from the abaxial side with a blunt end syringe. *N. benthamiana* leaves were imaged two days after infiltration.

### Confocal microscopy

GFP-tagged LYR4-ΔIC in *N. benthamiana* leaves was imaged using 488 nm excitation with 491 to 544 nm emission on a Zeiss LSM 780 confocal microscope.

### Expression and purification of kinases

*Medicago truncatula* LYR4 kinase (residues 298–660 or residues 340–636) and CERK1 (residues 307–598) were codon optimized for expression in *E. coli* (Genscript) and N-terminally fused with a histidine tag, an arginine tag, a SUMO tag, and a 3C protease cleavage site. LYR4 (residues 298– 660) A377F, LYR4 (residues 298–660) L392D, and CERK1 (307–598) K349A were made by site-directed mutagenesis of the aforementioned constructs. An overview of protein sequences used for constructs is provided in Additional Data 1. For expression, Rosetta 2 (Sigma-Aldrich) competent cells were transformed with the constructs and grown to OD_600_=0.6 in LB medium supplemented with the relevant antibiotics at 37°C with shaking. Then the culture was chilled on ice for 30 minutes before expression was induced by adding 0.2 mM IPTG and incubated at 19°C with shaking for 20 hours. The cells were harvested at 4,400 x g for 15 minutes, resuspended in LB medium, and pelleted again at 3,050 x g for 10 minutes. The pellet was resuspended in lysis buffer (50 mM Tris-HCl pH 8.0, 500 mM NaCl, 10% v/v glycerol, 10 mM imidazole, 5 mM β-mercaptoethanol (BME)), and lysed by sonication. Cleared supernatant was subjected to nickel-immobilized metal-affinity chromatography (Ni-IMAC) on a 1 mL Protino Ni-NTA column (Macherey-Nagel) equilibrated in lysis buffer. The sample was loaded onto the column using a peristaltic pump, then the column was washed with 20 column volumes buffer A (50 mM Tris-HCl pH 8.0, 500 mM NaCl, 10% v/v glycerol, 20 mM imidazole, 5 mM BME) before protein was eluted in 12 mL buffer B (50 mM Tris-HCl pH 8.0, 500 mM NaCl, 10% v/v glycerol, 200 mM imidazole, 5 mM BME). 3C protease was added to the eluate in an approximately 1:50 molar ratio and the digestion mixture was dialyzed against 1 L of dialysis buffer (50 mM Tris-HCl pH 8.0, 250 mM NaCl, 5% v/v glycerol, 5 mM BME) overnight at 4°C. The dialysis product was subjected to another Ni-IMAC to remove the protease and the digested tags before the protein was purified in a final gel filtration step on a Superdex75 increase 10/300 GL column or a Superdex200 increase 10/300 GL column (GE Healthcare) equilibrated in gel filtration buffer (25 mM Tris-HCl pH 8.0, 150 mM NaCl, 5 mM BME). All purification steps were analyzed by SDS-PAGE and elution fractions pooled accordingly.

### Crystallization and structure determination of the LYR4 kinase domain

The *Medicago truncatula* LYR4 kinase (residues 340–636) was crystallized in a sitting drop vapor diffusion setup in 24-well plates (Molecular Dimensions). Crystal formation in a condition with 0.2 M ammonium sulfate, 0.1 M sodium cacodylate trihydrate pH 6.5 and 30% w/v PEG 8000 gave rise to a seed stock used for microseeding of drops with protein and 4 mM adenylyl-imidodiphosphate (AMP-PNP) in a condition with 0.3 M ammonium sulfate, 0.1 M sodium cacodylate trihydrate pH 6.5, 25% w/v PEG 8000. Needle-like crystals were cryoprotected by supplementing the crystallization condition with 30% v/v glycerol before crystals were fished and flash-cooled in liquid nitrogen. The final data set was collected at Diamond Light Source beamline I04. The diffraction data was processed and scaled with XDS^31^, then analyzed in xtriage from the PHENIX program suite. Due to high anisotropy, data were scaled using the STARANISO server (Global Phasing Ltd. ^32^) along an anisotropic resolution limit surface (CC(1/2) >= 0.3, I/sigI >= 2.0, Rpim <= 0.6). The crystallographic phase problem was solved by molecular replacement using Phaser^33^. Refinement was performed in PHENIX ^34^ and manual model building was carried out in Coot^35^. Data collection statistics for both spherical and ellipsoidal datasets and refinement statistics are reported in Table S1. The atomic coordinates and structure factors were deposited in the Protein Data Bank, PDB ID: 8PS7. Structural analyses and figures were made using PyMol 2.3.2 (Schrödinger LLC).

### Differential scanning fluorimetry

Binding of the kinase to nucleotide was indirectly assayed with nano differential scanning fluorimetry (DSF) performed on a Tycho-NT.6 (Nanotemper). The experiments were made with LYR4 (residues 298–660), LYR4 A377F and LYR4 L392D at a 10 μM concentration in gel filtration buffer (25 mM Tris-HCl pH 8.0, 150 mM NaCl, 5 mM BME). ATP, AMP-PNP or MgCl_2_ were supplemented at a 1 mM concentration each.

### *In vitro* kinase assay

4 μg of each protein were incubated with 200 nCi [γ^32-P^] ATP (PerkinElmer) in 5 mM Tris-HCl pH 8.0, 30 mM NaCl, 5 mM MgCl_2_ and 20 mM cold ATP at room temperature for 1 hour. The samples were then separated on a 15% SDS-PAGE gel that was stained with InstantBlue (Expedion) before it was exposed overnight on an Autoradiography Hypercassette (Amersham/Cytiva). Radiographs were developed using a Typhoon FLA 9500 (Amersham/Cytiva).

