## Supplement for "The *Medicago truncatula* LYR4 intracellular domain serves as a scaffold in immunity signaling independent of its phosphorylation activity"

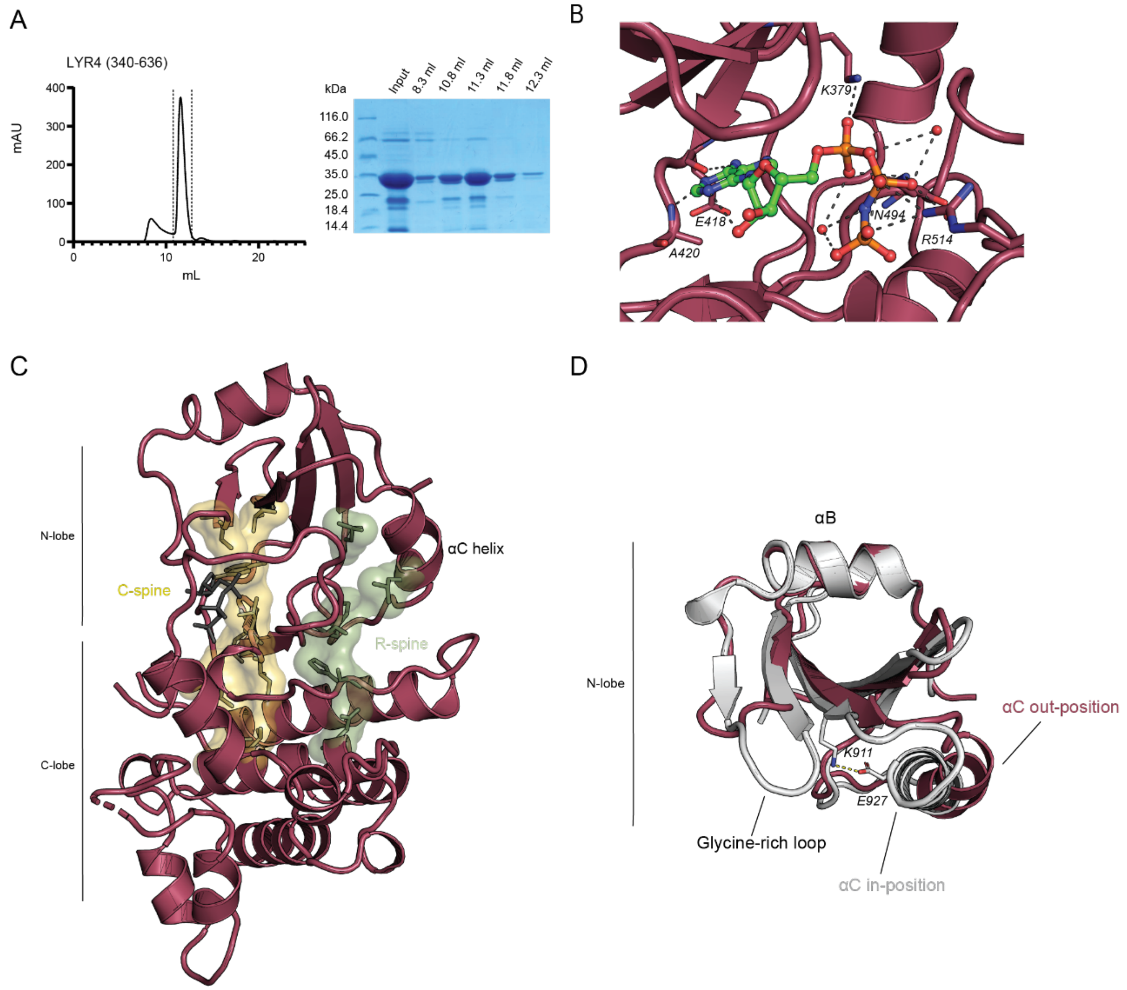


**Figure S1**. **Purification of LYR4 kinase domain and detailed structural analysis**. **A**. Gel filtration profile and Coomassie blue stained SDS-PAGE from purification of recombinant LYR4 (residues 340–636). Dashed lines indicate pooled fractions. **B**. Non-canonical AMP-PNP binding in LYR4 kinase. Dashed lines indicate molecular contacts between AMP-PNP and the kinase. Residues involved in nucleotide coordination are annotated and shown as sticks and water molecules involved in coordination are shown as red spheres. **C**. Crystal structure of LYR4 kinase shown with C-spine in yellow and R-spine in green. All residues that are part of the spines are shown as sticks. **D**. N-lobe of LYR4 kinase (shown in bordeaux) superimposed on the N-lobe of active kinase BRI1 (PDB-ID: 5LPV, shown in grey) to illustrate that LYR4 αC is in out-position. The salt bridge in BRI1 between regulatory lysine (K911) and conserved glutamate on the αC (E927) is shown with a dashed line.

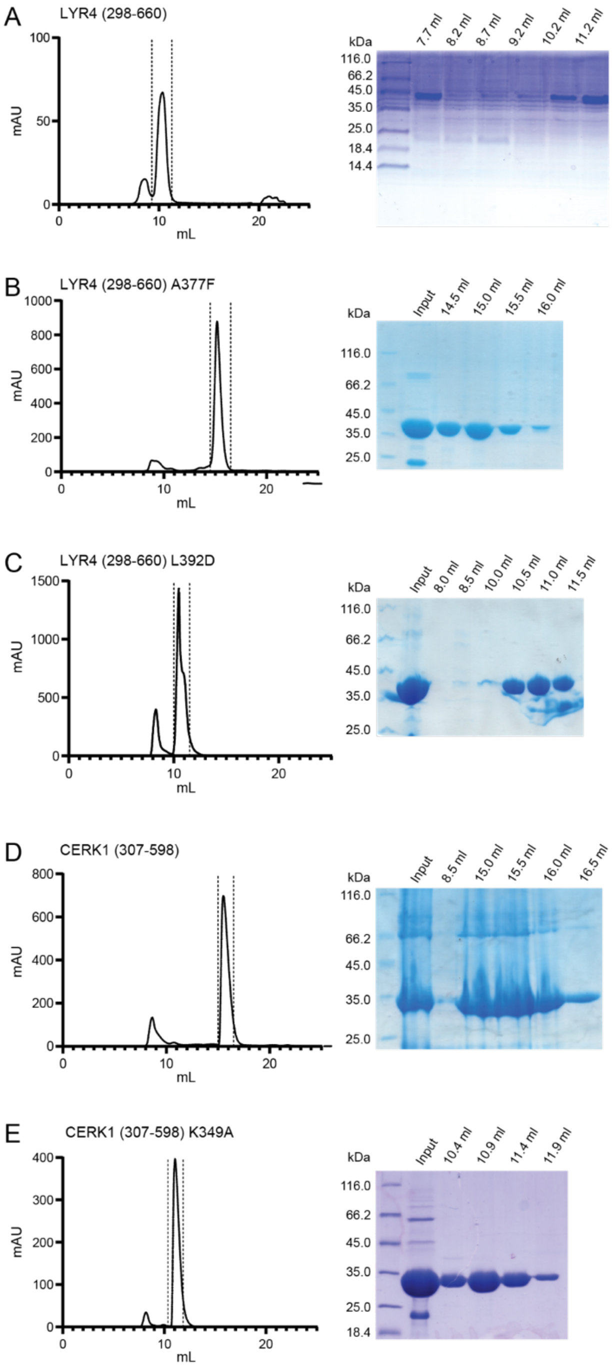


**Figure S2. Gel filtration profiles and corresponding SDS-PAGEs from purifications of recombinant proteins**. Dashed lines on chromatograms indicate pooled fractions. **A**. LYR4 (residues 298–660) purified on a Superdex75 increase 10/300 GL column. **B**. LYR4 (residues 298–660) A377F purified on a Superdex200 increase 10/300 column. **C.** LYR4 (residues 298–660) L392D purified on a Superdex75 increase 10/300 GL column. **D**. CERK1 (residues 307–598) purified on a Superdex200 increase 10/300 column. **F**. CERK1 (residues 307–598) K349A purified on a Superdex75 increase 10/300 GL column.

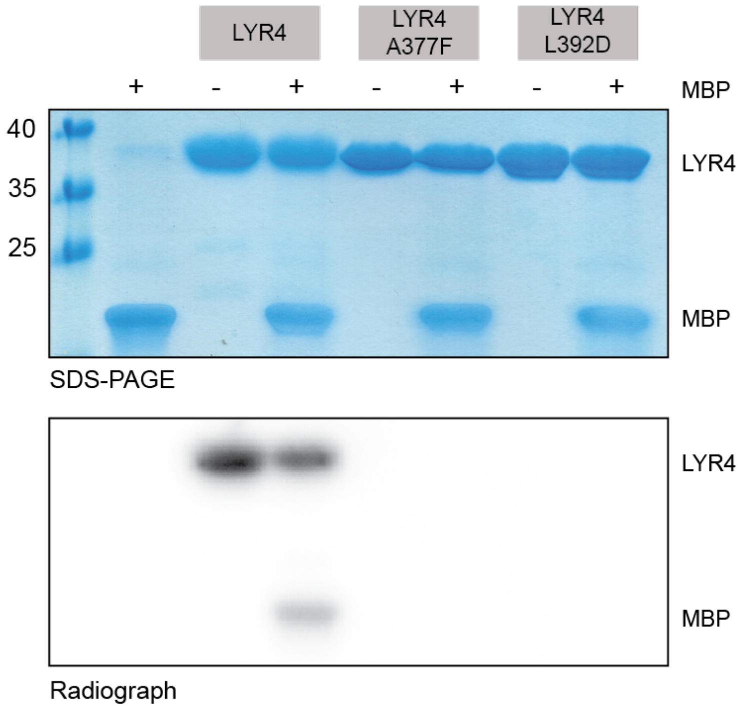


**Figure S3. *In vitro* kinase assay of LYR4 and LYR4 variants**. Intracellular domains of LYR4, LYR4-A377F and LYR4-L392D were incubated with radioactive ATP in the presence and absence of MBP as a transphosphorylation target. Top: Coomassie blue stained SDS-PAGE. Bottom: Corresponding radiograph.

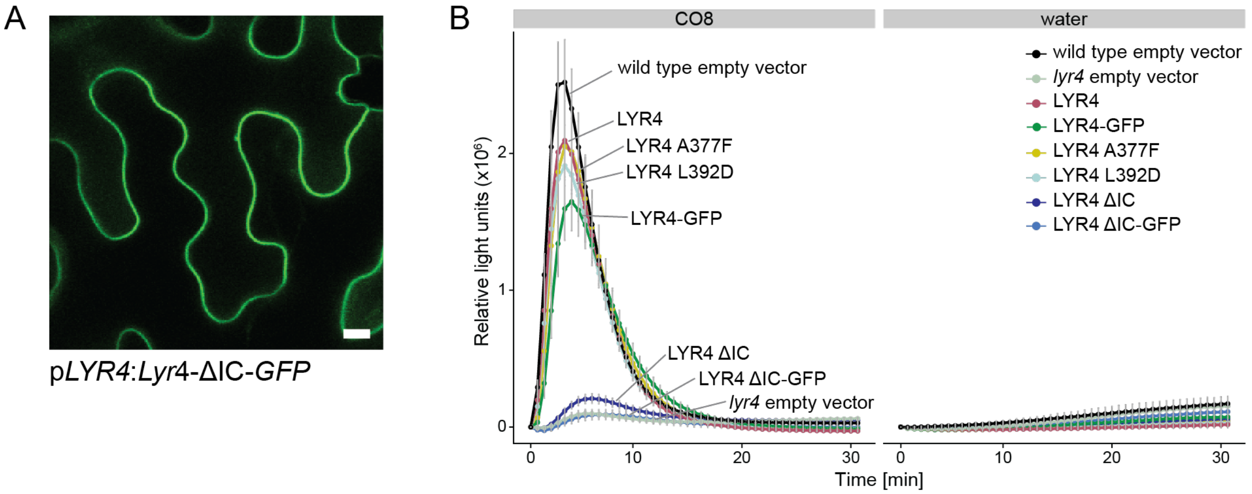


**Figure S4. ROS assays of LYR4 variants including GFP-tagged versions and membrane localization of LYR4-ΔIC**. **A**. Confocal image of GFP-tagged LYR4-ΔIC confirms production of the protein in *Nicotiana benthamiana* leaves. Scale bar is 10 µm. **B**. ROS production in *Medicago* wild type and *lyr4* single mutant roots transformed with *Lyr4*, *Lyr4-GFP*, *Lyr4*-A377F, *Lyr4*-L392D, *Lyr4*-ΔIC or *Lyr4*-ΔIC-*GFP* over a period of 30 minutes after treatment with 1 µM CO8 or water (mean ± standard error of the mean, n = 15 (CO8) or n = 3 (water)). The data shown are the same as in Figure 3C but include GFP-tagged variants.

### Table 1 – Data collection and refinement statistics

| Data collection | Elliptical trunc. | Spherical trunc. |
|---|---|---|
| **X-ray source** | DLS I04 | DLS I04 |
| **Wavelength (Å)** | 0.97 | 0.97 |
| **Space group** | $P12_11$ | $P12_11$ |
| ***Unit cell parameters*** | | |
| **a, b, c (Å)** | 51.68, 78.24, 70.36 | 51.68, 78.24, 70.36 |
| **Α, β, γ (°)** | 90.0, 90.4, 90.0 | 90.0, 90.4, 90.0 |
| **Resolution (Å)** [a] | 43.1 – 2.1 (2.3 – 2.1) | 43 – 3.1 (3.2 – 3.1) |
| **Upper limit along a*,b*c* (Å)** | 2.5, 3.0, 2.1 | |
| **Unique reflections** [a] | 18580 (929) | 10270 (924) |
| **Redundancy** [a] | 3.5 (3.5) | 3.5 (3.4) |
| **Completeness (%)** [a] | 91.2 (70.2) | 99.7 (99.7) |
| **$R_{meas}$ (I) (%)** [a,] | 24.4 (90.3) | 15.1 (45.5) |
| **I/σ (I)** [a] | 5.2 (1.6) | 8.6 (3.1) |
| **CC(1/2)** [a] | 98.1 (62.0) | 99.0 (88.7) |
| **Wilson B** | 21.79 | 31.9 |
| **Refinement statistics** | | |
| **Resolution (Å)** [a] | 43.1-2.1 (2.3-2.1) | |
| **Reflections ($R_{work}$)** | 17113 | |
| **Reflections ($R_{free}$)** | 1464 | |
| **$R_{work}$ (%)** | 22.1 | |
| **$R_{free}$ (%)** | 26.9 | |
| **No. of atoms in a.u.** | | |
| **protein** | 4383 | |
| **ligands and solutes** | 124 | |
| **water** | 173 | |
| **R.M.S. deviations** | | |
| **bonds (Å)** | 0.002 | |
| **angles (°)** | 0.5 | |
| ***Average B values*** | | |
| **protein (Å²)** | 35.5 | |
| **ligands (Å²)** | 50.0 | |
| **water (Å²)** | 31.1 | |
| ***Ramachandran plot*** | | |
| **favored (%)** | 94.8 | |
| **allowed (%)** | 5.2 | |
| **outliers (%)** | 0 | |

[a] Values for the highest resolution shell are given in parentheses.
